# Machine Learning-Based Early Prediction of Neural Microelectrode Lifetime Under Accelerated Aging Conditions

**DOI:** 10.64898/2026.10.29.762829

**Authors:** Hui Hua, Martin Niemiec, Tingkai Li, Rahul Manna, Hyeongjin Jo, Kyungjin Kim, Chao Hu

## Abstract

**Objective:** Chronically implantable medical devices may require functional lifetimes of decades in order to reduce or eliminate the need for their explantation and replacement, which may involve invasive and sometimes risky surgery. However, when devices become exceptionally long-lived, preclinical studies validating their lifetimes become difficult to execute within reasonable timeframes even when accelerated life testing methods are employed. Computational methods capable of accurately and reliably predicting microelectrode lifetime from early-life data could therefore considerably expedite the development of long-lived implantable devices.

**Approach:** Here, inspired by similar studies predicting functional lifetimes of lithium-ion battery cells from early-life data, we developed a machine learning model based on Elastic Net regression and early-life feature engineering to predict impedance-defined microelectrode lifetimes under accelerated aging conditions and evaluated its predictive performance. We also provide a dataset of electrochemical measurements from aging microelectrodes for public use.

**Main results:** Using measurements collected through Day 5, the 10-feature Elastic Net model achieved a root mean squared error of 3.71 accelerated-aging days and a mean absolute percentage error of 13.90% under device-level validation. In an independent fabrication batch, covariance alignment and calibration using five labeled channels from one device in each target group reduced mean absolute percentage errors from 38.1% to 22.5% for Small electrodes and from 56.6% to 18.2% for Medium electrodes.

**Significance:** These results highlight the potential of machine learning-based approaches to accurately predict neural microelectrode lifetimes from early-life data. Such methods may inform the design of future accelerated life studies, providing meaningful longevity estimates from shorter testing periods and potentially accelerating the development of long-lived implantable devices for clinical use.

## 1 Introduction

The long-term functional stability of implantable medical devices is critical for chronic applications. [1, 2] Neural interfaces encompass diverse device architectures, including rigid silicon-based Utah electrode arrays, flexible polymer-based microelectrode arrays (MEAs), and soft elastomeric interfaces. [3, 4, 5] These platforms differ in their mechanical properties, electrode configurations, and material interfaces, creating different requirements for long-term reliability. In particular, device lifetime can vary substantially depending on device design and configuration, as well as the materials and thin-film encapsulation strategies used to protect metal tracks against degradation in physiological environments. [6] These strategies include vacuum-deposited inorganic dielectric films, such as silicon carbide prepared by plasma-enhanced chemical vapor deposition (PECVD) and metal oxides prepared by atomic layer deposition (ALD), used alone or combined with polymer layers. [7, 8, 9, 10] Atomic layer infiltration (ALI) provides a related approach in which inorganic species are introduced into the polymer insulation. [11] Differences in device geometry, layer architecture, material interfaces, and encapsulation design can influence degradation mechanisms and ultimately determine functional lifetime. [8, 12] Beyond encapsulation-related failure, localized pitting corrosion of thin-film microelectrodes under electrical bias in chloride-containing environments can cause material loss and compromise long-term reliability. [13] To evaluate device functionality and degradation, in vitro soaking tests in phosphate-buffered saline (PBS) are commonly performed, while electrochemical impedance spectroscopy (EIS) and cyclic voltammetry (CV) are used to monitor changes in electrochemical performance over time. [14, 15]

Chronically implanted devices, particularly neural interfaces, may require functional lifetimes on the order of decades to minimize the need for device explantation and replacement, which can involve additional invasive surgical procedures and risks to surrounding tissues. [4] However, validating such long functional lifetimes through conventional aging studies is challenging within practical experimental time frames. Accelerated life testing at elevated temperatures is therefore commonly employed to accelerate degradation and estimate device lifetime under physiological conditions. [16] For example, accelerated aging studies of electrohydrodynamically printed transparent soft microelectrodes have been used to assess electrochemical stability and project multi-year operational lifetimes. [17] Reactive accelerated aging using reactive oxygen species has also been developed for rapid evaluation of cortical neural implant durability. [18] Comparisons between reactive accelerated aging and multi-year in vivo implantation have been used to assess the relevance of laboratory-observed Parylene-C degradation mechanisms. [19] Nevertheless, highly stable devices may still require several weeks to months or longer of aging and periodic electrochemical monitoring before reaching defined failure criteria. This limits experimental throughput and makes rapid comparison of different device designs, materials, and encapsulation strategies difficult. [20]

Machine learning-based lifetime prediction provides a potential approach to overcome this limitation. Similar approaches have been investigated for lithium-ion batteries, where features extracted from early cycling behavior are used to predict eventual battery lifetime without waiting for each new battery to reach its end of life. [21, 22]

Related approaches combine empirical capacity-fade models with machine learning to predict battery degradation trajectories from early-life features. [23] Hybrid data-driven and model-based methods have also been investigated for estimating the remaining useful life of lithium-ion batteries in implantable medical devices. [24] This concept can potentially be extended to neural microelectrodes, as electrochemical measurements collected during the early stages of accelerated aging may contain information associated with progressive degradation and eventual device failure.

Herein, a machine learning-based approach for predicting microelectrode lifetime using early-life electrochemical data collected under accelerated aging conditions is presented. The present study focuses on platinum microelectrodes on flexible polyimide substrates, including devices with polyimide insulation alone (Control) and devices with additional ALD or ALI-based inorganic encapsulation. This device and material system provides a defined setting for evaluating early-life lifetime prediction across electrode sizes, encapsulation conditions, and fabrication batches. An Elastic Net regression model combined with early-life feature engineering is used to establish relationships between early aging behavior and eventual device lifetime. By integrating accelerated aging experiments with data-driven lifetime prediction, this approach may substantially reduce the testing time needed to estimate microelectrode lifetime and enable more efficient evaluation of reliable and long-lived implantable neural interfaces.

## 2 Dataset and Problem Formulation

### 2.1 Neural microelectrode aging dataset

The dataset used to train and validate the lifetime prediction model consisted of electrochemical measurements from platinum microelectrodes on polyimide substrates, specifically electrochemical impedance spectroscopy (EIS) and cyclic voltammetry (CV) data. The dataset primarily consisted of data originally collected for a previous publication [11], in which details of the device fabrication and test parameters can be found. Briefly, platinum microelectrodes of geometric surface area (GSA) 1,000*µ*m^2^ (small), 10,000*µ*m^2^ (medium), and 100,000*µ*m^2^ (large) were fabricated on thin-film polyimide substrates (total thickness appx. 7 *µ*m) and aged in phosphate buffered saline (PBS) at 87^*°*^C. Each day, EIS spectra were collected from 0.1Hz–100kHz and cyclic voltammetry was performed, sweeping between −0.6V and 0.8V at a rate of 100mV/s. Tests were performed in PBS maintained at 37^*°*^C in a water bath, to reflect the temperature that would be experienced by implanted devices.

To increase the sample size, additional devices were fabricated for this work following the fabrication steps described in [11] as closely as possible so that the failure modes of the newly fabricated devices matched those previously reported. The original dataset consisted of 77 channels across 17 devices, with a total of 31 small, 28 medium, and 18 large sites analyzed. 15 channels of each size were added to the dataset after the new devices were fabricated and tested, bringing the total to 46 small, 43 medium, and 33 large sites. Some parameters of the newly tested devices were varied from the previous study in order to reduce total test duration per channel, enabling more devices to be tested per day. For example, the number of EIS samples per decade frequency was decreased from 12 to 6 for the first week of aging, upon which predictions were based, after which point sampling was reduced to 3 points per decade, reducing test duration while still providing enough information to identify and characterize failure. The full dataset is openly available in Zenodo (see Data availability).

### 2.2 Failure criterion and lifetime definition

In the original work [11], failure was defined in two ways: one purely by change in impedance, where any channel deviating in impedance by more than 50% in either direction from its original value after stabilization was considered to have failed, and the other more subjectively based on synthesis of additional sources of information such as optical images. For the purposes of machine learning-based prediction, only the impedance-based definition of failure has been utilized in this work. Specifically, for each tested channel *i*, the failure day *t*_*f,i*_ was defined as the first day on which the relative change in 1 kHz impedance exceeded 50% from the stabilized initial impedance:

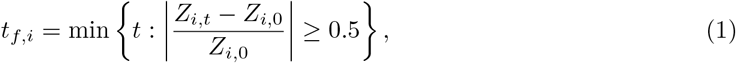

where *Z*_*i*,0_ is the stabilized initial 1 kHz impedance of channel *i*, and *Z*_*i,t*_ is the 1 kHz impedance measured on day *t*.

This metric is somewhat arbitrary, as in the absence of true functional evaluations which might be performed *in vivo* (such as ability to evoke action potentials or signal-to-noise ratio (SNR) of recorded neural activity), electrode health and functionality are more difficult to define *in vitro*. It is also worth observing that impedance changes associated with different failure modes may offset each other.

For example, delamination of insulation will typically lead to decreased impedance, while a crack developing in an electrode’s conductive trace will tend to increase impedance, and both of these failure modes can occur simultaneously. Nonetheless, impedance deviation provides a measurable and objective metric that, in vivo, generally correlates with a channel’s ability to perform the aforementioned functions, and was therefore considered a feasible criterion of failure.

The lifetime *L*_*i*_ of channel *i* was defined as the elapsed aging time from the initial baseline measurement to the impedance-based failure day:

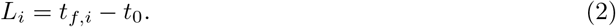

Because the baseline measurement was defined as Day 0 in this study, *t*_0_ = 0, and therefore:

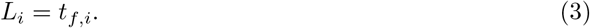

### 2.3 Early-life prediction problem formulation

The early-life prediction task was formulated at the channel level. Each tested channel was treated as one prediction sample, with its early electrochemical response being used to predict its lifetime under accelerated aging conditions. Let Day *k* denote the early prediction time point, where *k* ≤ 5. Only measurements from Day 0 to Day *k* were used for feature extraction, and no information collected after Day *k* was included in the model input.

For channel *i*, the engineered early-life feature vector constructed using data available up to Day *k* is denoted as 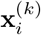. This feature vector contains fixed-day features extracted from Day *k* measurements and cross-day features calculated relative to the Day 0 baseline. The prediction target is the lifetime *L*_*i*_, which is equivalent to the failure day *t*_*f,i*_ defined in Sec. 2.2. Therefore, the experimental dataset for the Day *k* prediction task can be written as

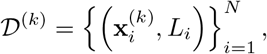

where *N* is the number of tested channels for which measurements are available from Day 0 to Day *k* and a failure has been observed.

The objective is to learn a regression function *f* ^(*k*)^(·) that maps the early-life feature vector to the predicted lifetime:

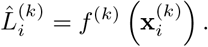

The prediction error is then evaluated by comparing the predicted lifetime 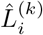 with the observed lifetime *L*_*i*_. This formulation allows the model to predict the final lifetime using only early-life electrochemical information.

## 3 Early-Life Feature Engineering

### 3.1 Overview of Early-Life Feature Construction

Our approach estimates each microelectrode channel’s lifetime from electrochemical measurements collected during the early stages of accelerated aging. Therefore, feature engineering is needed to extract informative descriptors from limited early-life data that can reflect both the device’s state and its degradation trend. Based on this motivation, we developed an early-life feature engineering pipeline to extract complementary degradation information from early EIS and CV data.

For each tested channel, two types of early-life features were constructed: fixed-day features and cross-day features. The fixed-day features were extracted from measurements at *Day k*, where (*k* ≤ 5), aiming to describe the electrochemical state of the device at the early prediction time point. The cross-day features were calculated based on the changes in electrochemical measurements relative to the *Day* 0 baseline, which characterize the evolution of the internal state from the early-stage aging.

As mentioned, the overall feature pool consisted of two source-based groups: EIS-based features and CV-based features. Each group also included raw statistical features and derived features. As summarized in Table 1, the resulting candidate feature pool contained 2025 features in total. The construction of each feature group is described in the following sections, with the complete feature definitions and extraction details provided in the Supplementary Material.

**Table 1.** Summary of the engineered early-life feature pool. Fixed-day features are extracted at *Day k*; cross-day features are calculated relative to the *Day* 0 baseline.

| Modality | Feature type | Fixed-day features | Cross-day features | Total |
| --- | --- | --- | --- | --- |
| EIS | Raw | 160 | 240 | 400 |
| EIS | Derived | 528 | 543 | 1071 |
| CV | Raw | 127 | 127 | 254 |
| CV | Derived | 150 | 150 | 300 |
| Total | – | 965 | 1060 | 2025 |

### 3.2 EIS-based features

#### 3.2.1 Raw EIS features

Raw EIS features were extracted from the impedance magnitude and phase spectra. For each tested channel, the EIS measurements were first processed to obtain frequency, real impedance, imaginary impedance, impedance magnitude, and phase. The spectra were sorted by frequency, and the impedance magnitude and phase were interpolated on the logarithmic frequency axis to ten fixed frequencies: 0.1, 0.3, 1, 3, 10, 30, 100, 1,000, 10,000, and 100,000 Hz. These fixed-frequency features preserve the impedance response at representative points across the measured frequency range. In addition to fixed-frequency values, global statistical descriptors were extracted from the impedance magnitude and phase spectra, including the mean, standard deviation, variance, minimum, maximum, median, range, skewness, excess kurtosis. The area under each curve with respect to logarithmic frequency was also calculated.

Frequency-band-based features were further extracted to capture local impedance behavior within different frequency regions. The predefined frequency bands included both adjacent bands in the logarithmic scale and wider overlapping bands: 0.1–1 Hz, 1–10 Hz, 10–100 Hz, 100–1,000 Hz, 1,000–10,000 Hz, 10,000–100,000 Hz, 0.1–10 Hz, 1–100 Hz, 10–1,000 Hz, and 100–10,000 Hz. For each frequency band, the mean, standard deviation, minimum, maximum, slope, and area with respect to logarithmic frequency were computed. Together, these raw EIS statistical features summarize the overall impedance magnitude, phase distribution, and frequency-dependent response of the microelectrodes during early aging.

#### 3.2.2 Derived EIS features

The derived EIS features were designed to capture higher-level shape information from the impedance response. First, derivative-based features were calculated from the full EIS spectrum with respect to logarithmic frequency. Five derivative curves were computed, including the derivatives of real impedance, imaginary impedance, impedance magnitude, phase, and logarithmic impedance magnitude with respect to log_10_(*f*). From each derivative curve, global statistical descriptors, extreme-value features, and frequency-band-based features were extracted. The extreme-value features included the maximum and minimum derivative values, their corresponding frequencies, peak-to-valley amplitude, peak-to-valley distance on the logarithmic frequency axis, and absolute area. These derivative-based features were designed to characterize local changes and transition patterns in the impedance spectrum.

Geometric features on the Nyquist plot were also extracted from the real and negative imaginary impedance responses. These features included high- and low-frequency geometric anchors, the maximum negative imaginary impedance and its corresponding real impedance and frequency, the span of real and negative imaginary impedance, Nyquist area, arc height, arc width, arc aspect ratio, low-frequency tail slope, and the low-frequency ratio between negative imaginary impedance and real impedance. Compared to raw statistical descriptors, these geometry-based features directly characterize the shape of the Nyquist plot and may capture electrode-interface characteristics associated with early degradation.

### 3.3 CV-based features

#### 3.3.1 Raw CV features

Raw CV features were extracted from the current-potential curves to characterize the early electrochemical response of each microelectrode. For each tested channel, the CV data were processed to obtain potential and current measurements. To ensure a consistent representation of the CV response, the last 2,800 data points from each CV measurement were used for feature extraction. Similar to raw EIS features, global statistical descriptors were calculated from the current signal, as well as the signed area and absolute area with respect to potential. In addition, positive and negative current areas and the fractions of positive and negative current values were extracted to describe the polarity distribution of the CV response.

Extreme-value features were also extracted from the current-potential curve, including the maximum and minimum current values, their corresponding potentials, the peak-to-valley amplitude, and the potential distance between the peak and valley. To capture voltage-dependent behaviors, window-based features were computed over predefined potential ranges. Within each voltage window, statistical and shape descriptors, including the mean, standard deviation, minimum, maximum, median, range, area, absolute area, and slope, were extracted. Together, these raw CV statistical features characterize the current magnitude, polarity distribution, extrema, and voltage-dependent response of the microelectrodes during early aging.

#### 3.3.2 Derived CV features

Derived CV features were extracted from the derivative response of the current-potential curve. Before calculating the derivative, the current signal was smoothed using a Savitzky–Golay filter. The derivative curve was then computed as *dI/dV* using the ratio between the numerical gradient of current and the numerical gradient of potential. Data points with near-zero potential gradients were excluded to avoid unstable derivative values.

From each *dI/dV* curve, global statistical descriptors were extracted, including the mean, standard deviation, variance, minimum, maximum, median, range, skewness, excess kurtosis, area, and absolute area. Extreme-value features were also computed, including the maximum and minimum *dI/dV* values, their corresponding potentials, the peak-to-valley amplitude, and the potential distance between the peak and valley. In addition, voltage-window-based *dI/dV* features were extracted to capture localized changes in the CV curve shape. These derived CV features provide complementary information to the raw current descriptors by emphasizing local changes in the current-potential relationship.

## 4 Methodology

### 4.1 Overview of the proposed framework

Figure 1 provides an overview of the proposed framework for early-life prediction of neural microelectrode lifetime. The goal is to predict the final lifetime using limited measurements collected during the early stages of accelerated aging, without waiting for each new device to reach failure. The framework consists of four main stages: electrochemical aging data collection, early-life feature engineering, feature selection and model development, and early-life lifetime prediction.

**Figure 1.**
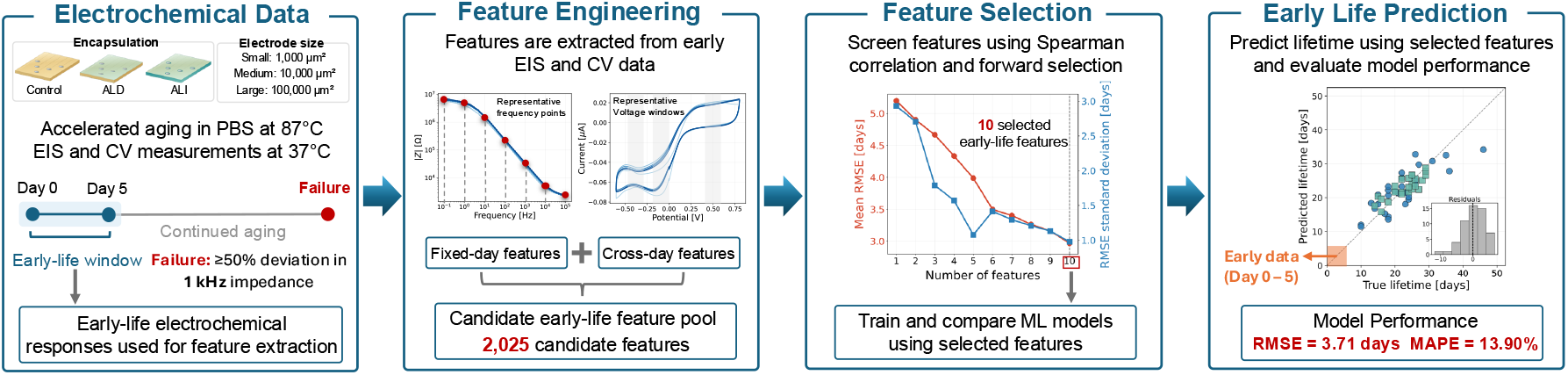
Overview of the proposed framework for early-life prediction of neural microelectrode lifetime.

Although the electrochemical aging dataset covers the full aging process, only EIS and CV measurements from the early-life window were used for feature engineering. From these measurements, we extracted a large pool of candidate features describing the early electrochemical state and its changes over time. Feature screening and selection were then performed to identify informative feature combinations, which were used as inputs to machine learning regression models. The trained models were used to predict the lifetime of each microelectrode channel, and prediction performance was evaluated by comparing predicted and observed lifetimes. In addition, the framework was evaluated using data from an independently fabricated batch to assess its predictive performance under distribution shift.

### 4.2 Partitioning data for machine learning

In this study, the primary modeling dataset consisted of Small and Medium electrode sites. Large sites were excluded from model development and evaluation because our previous study found greater susceptibility to metal cracking at the electrode–trace junction in these sites. This behavior was attributed to stress concentration at the abrupt transition from the narrow conductive trace to the wider electrode and was associated with shorter lifetimes [11].

Of the original 59 Small and Medium channels in Batch 1, four exhibiting premature failure associated with bond pad cracking (6–11 days) were excluded from the modeling analysis. As summarized in Table 2, the remaining Small and Medium electrodes from Batch 1 formed the model-development dataset, consisting of 13 devices and 55 channels. The Small and Medium electrodes from Batch 2 were treated as two separate out-of-distribution test sets, each containing 3 devices and 15 channels.

**Table 2.**
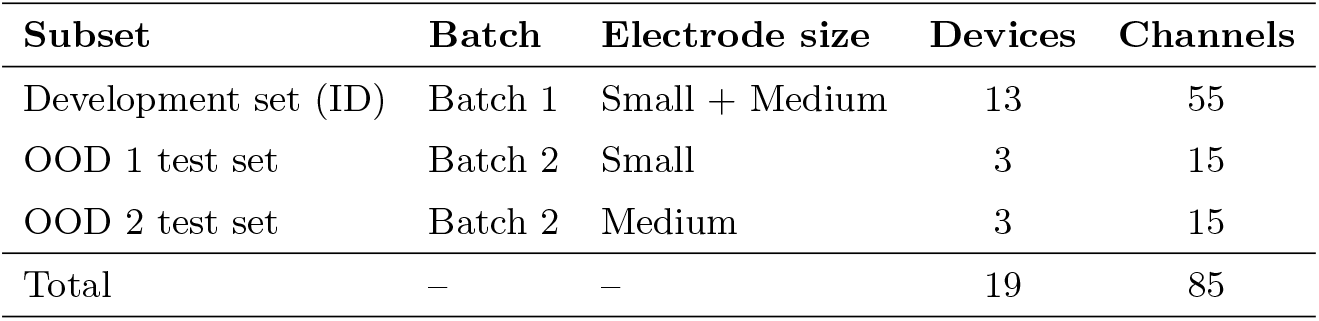
Summary of data partitioning for model development and out-of-distribution evaluation.

| Subset | Batch | Electrode size | Devices | Channels |
| --- | --- | --- | --- | --- |
| Development set (ID) | Batch 1 | Small + Medium | 13 | 55 |
| OOD 1 test set | Batch 2 | Small | 3 | 15 |
| OOD 2 test set | Batch 2 | Medium | 3 | 15 |
| Total | — | — | 19 | 85 |

Batch 2 consisted of independently fabricated and tested devices and exhibited longer lifetimes than Batch 1. As shown in Fig. 2, the increase was substantially greater for Medium than for Small electrodes. Delamination and cracking were observed in both batches, although cracking may have contributed more strongly to failures in Batch 1. More frequent handling for daily imaging in Batch 1 may have promoted crack initiation and propagation, potentially limiting lifetimes in both Small and Medium electrodes. In contrast, less frequent handling in Batch 2 may have reduced the contribution of cracking, particularly for Medium electrodes. Although cracks were still observed, those that had not propagated across the conductive trace may not have caused immediate loss of electrical continuity.

**Figure 2.**
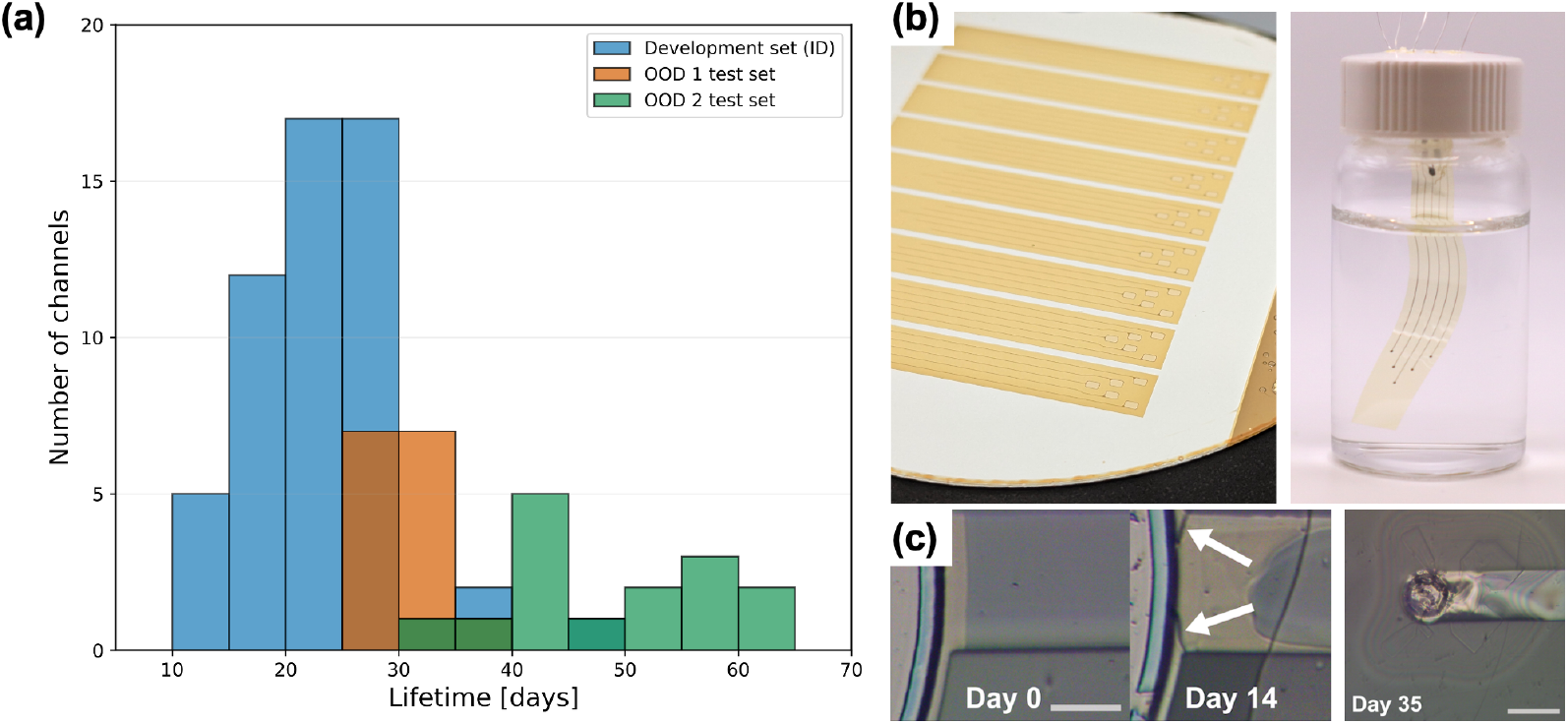
Channel lifetime distributions, device photographs, and representative failure modes of polyimide microelectrode arrays (MEAs). (a) Distribution of channel lifetimes in the Batch 1 development set (ID; Small and Medium electrodes), the Batch 2 Small test set (OOD 1), and the Batch 2 Medium test set (OOD 2). (b) Photographs of fabricated polyimide MEAs on a wafer (left) and a released MEA immersed in phosphate-buffered saline (PBS) in the soaking setup used for accelerated aging (right). (c) Representative optical micrographs showing an electrode–trace junction before (Day 0) and after cracking (Day 14; white arrows indicate cracks), alongside a separate example of delamination between polyimide layers and between polyimide and metal traces (Day 35). Scale bars in (c): 25 *µ*m (Day 0 and Day 14) and 50 *µ*m (Day 35). Images in panel (c) adapted from [11], © 2025 the authors, under the CC BY licence.

For Small electrodes, delamination around the electrode opening appeared to be an important failure mechanism in both batches, potentially explaining their comparatively modest between-batch lifetime difference. Although Batch 2 Medium electrodes also exhibited delamination, their larger initial active electrode area may have reduced the relative impedance change associated with a comparable amount of additional conductive area exposed by localized delamination. Together with reduced cracking, this may have delayed reaching the 50% impedance-change threshold used to define failure. These explanations remain qualitative, as the individual contributions of handling and electrode geometry were not independently isolated.

Small and Medium electrodes from Batch 1 were pooled to develop a common prediction model spanning both electrode sizes. The Batch 2 groups were evaluated separately because their distinct lifetime distributions, particularly the substantially longer lifetimes of Medium electrodes, allowed prediction performance to be assessed under different degrees of distribution shift.

Although lifetime was defined at the channel level, all data partitioning and cross-validation were performed at the device level. Each polymer microelectrode array contains multiple electrode channels, and channels from the same device share the same fabrication process, coating condition, and aging environment. Therefore, all channels from a given device were kept within the same data subset to avoid information leakage.

### 4.3 Stepwise Forward Feature Selection

The feature pool contains EIS- and CV-based descriptors, which may be correlated with each other and provide complementary predictive information. Therefore, a multivariate feature selection strategy was used to identify informative feature combinations for lifetime prediction.

To reduce model complexity and the risks of overfitting, we performed stepwise forward feature selection with a linear model. The stepwise forward selection procedure started with an empty feature set. At each step, each remaining candidate feature was added individually to the current selected feature set, and the linear model was evaluated using five-fold device-level cross-validation. The feature that achieved the lowest mean cross-validation root mean squared error (RMSE) was added to the selected subset. This process was repeated until up to 10 features were selected. The final feature subset was determined by considering both the mean and standard deviation of the cross-validation RMSE, favoring subsets with low prediction error and stable performance across folds. Feature selection was performed once using the full Batch 1 development dataset. The resulting 10-feature subset was held fixed across all outer cross-validation folds and repetitions for subsequent model comparisons. Batch 2 data were not used for feature selection. From a practical application perspective, smaller feature subsets were preferred to support efficient inference.

### 4.4 Machine Learning Models for Lifetime Prediction

#### 4.4.1 Elastic Net Regression

Elastic Net regression was used as the primary model for lifetime prediction. It is a regularized linear regression model that combines the *L*_1_ penalty used in LASSO regression and the *L*_2_ penalty used in Ridge regression.

For each tested channel *i*, the selected early-life feature vector is denoted as **x**_*i*_ ∈ ℝ^*p*^, and the corresponding observed lifetime is denoted as *L*_*i*_. The Elastic Net model predicts lifetime as

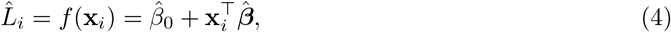

where 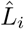 is the predicted lifetime, 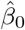 is the fitted intercept, and 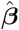 is the fitted vector of regression coefficients. The model coefficients were estimated by minimizing the following regularized objective function:

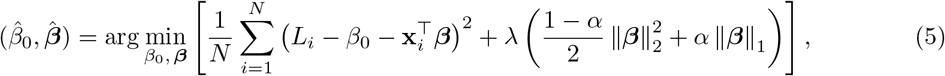

where *N* is the number of training samples, *λ* controls the overall strength of regularization, and *α* controls the balance between the *L*_1_ and *L*_2_ penalties. The regularization parameters *λ* and *α* were selected by cross-validation on the training set.

#### 4.4.2 Comparison Models

To evaluate the predictive performance of the selected early-life features across different machine learning models, we included Ridge regression, Gaussian process regression (GPR), and LightGBM for comparison.

**Ridge regression** is a linear model with *L*_2_ regularization that penalizes large coefficients and helps reduce overfitting. Unlike Elastic Net, Ridge does not include an *L*_1_ penalty, allowing comparison of different regularization strategies within linear models.

**GPR** is a kernel-based probabilistic regression method that can capture nonlinear relationships between the input features and lifetime. Three candidate kernels, including RBF and Matérn (*ν* = 1.5 and 2.5), were evaluated, with the optimal kernel selected through cross-validation. GPR was included to assess nonlinear regression performance with limited training data.

**LightGBM** is a gradient-boosted decision tree model that builds an ensemble of trees iteratively to reduce prediction error. It can capture nonlinear relationships and feature interactions, and was included to evaluate whether a more flexible tree-based model could improve lifetime prediction.

All comparison models used the same feature subset and device-level validation strategy as Elastic Net to ensure a consistent comparison. Detailed model configurations and hyperparameter search spaces are provided in the Supplementary Material.

### 4.5 Performance evaluation metrics

The lifetime prediction performance was evaluated using two metrics: RMSE and mean absolute percentage error (MAPE). These metrics are defined as follows:

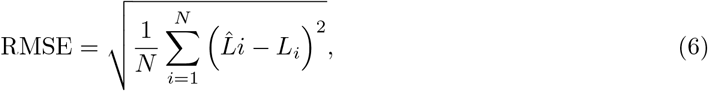

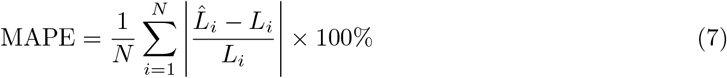

where *N* is the number of evaluated samples, 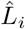 is the predicted lifetime of the *i*-th tested channel, and *L*_*i*_ is the corresponding observed lifetime. RMSE measures the absolute prediction error in days, whereas MAPE quantifies the relative prediction error as a percentage.

## 5 Results and discussion

### 5.1 Feature Screening and Forward Feature Selection

The initial early-life feature pool contained 2,025 candidate features derived from EIS and CV measurements. Because many features were extracted using densely spaced and overlapping frequency or voltage windows, substantial correlation and redundancy were likely to exist within the feature pool. To reduce this redundancy before multivariable feature selection, three screening strategies were compared, as shown in Fig. 3. For redundancy filtering, features with an absolute pairwise Spearman correlation of |*ρ*| ≥ 0.85 were considered highly redundant.

**Figure 3.**
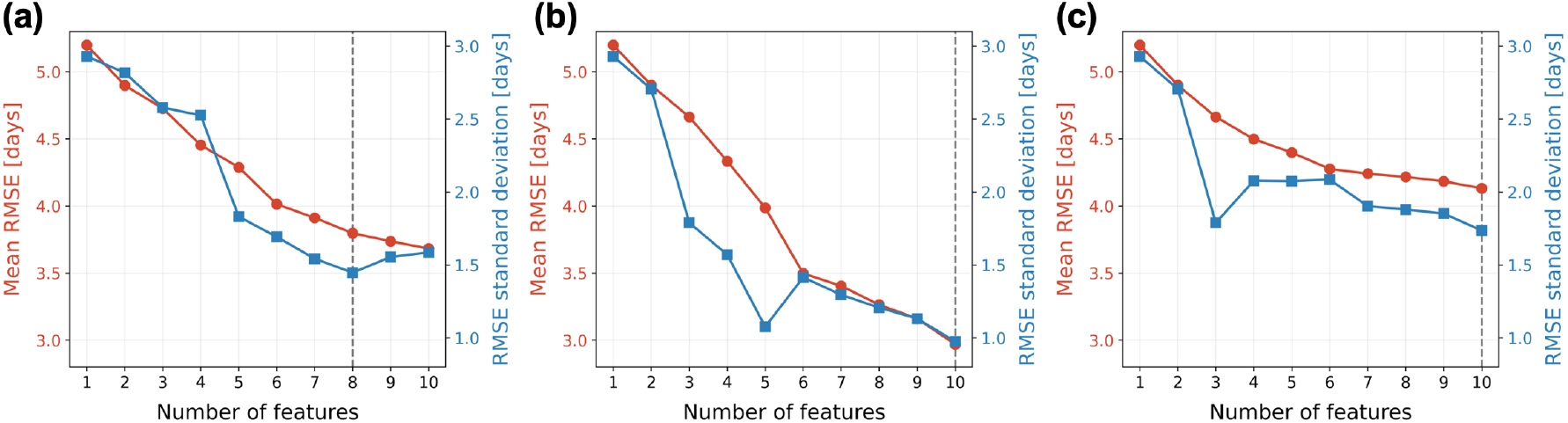
Forward feature selection results for three feature-screening strategies: (a) the full 2,025-feature pool, (b) the redundancy-filtered feature pool, and (c) the redundancy-filtered Top-100 feature pool. The gray dashed line marks the selected number of features, determined by minimizing the sum of the mean CV RMSE and its standard deviation.

Redundancy filtering reduced the candidate pool from 2,025 to 370 features. Among the three strategies, the redundancy-filtered feature pool achieved the lowest device-level cross-validation RMSE. Because many features were extracted from adjacent or overlapping frequency and voltage windows, some candidate features likely captured similar electrochemical information. Retaining all 2,025 features substantially increased the search space without necessarily adding new predictive information. With the limited sample size, this may also make forward selection more sensitive to the composition of the development data. Removing highly correlated features therefore reduced the search space while preserving more distinct sources of information, leading to a more efficient and stable selection process.

In contrast, further restricting the redundancy-filtered pool to the 100 features with the strongest Spearman correlations with lifetime degraded the prediction performance. This suggests that features with moderate or even weak individual correlations with lifetime may still provide useful complementary information when combined with other predictors.

For the redundancy-filtered feature pool, the CV RMSE decreased consistently as additional features were selected, with a particularly noticeable reduction between approximately three and six features. This indicates that the added features provided additional non-redundant information. In comparison, the Top-100 feature pool reached a plateau earlier, suggesting more limited gains from adding further features. Given the limited sample size, the forward search was restricted to a maximum of 10 features. Therefore, the 10-feature subset represents the best result within the current search range, rather than a confirmed global optimum.

### 5.2 Selected Early-Life Lifetime Features

To control model complexity and reduce the risk of overfitting, 10 features were retained for subsequent analysis. Table 3 summarizes the 10 features added sequentially during stepwise forward feature selection. The final subset included eight EIS-based features and two CV-based features, covering both state and change features. Fig. 4(a–d) illustrates how the first two selected features were extracted from early-life electrochemical measurements: the CV positive-current fraction and the change in the EIS phase derivative. Fig. 4(e) presents the correlation matrix of the 10 selected features. Overall, most feature pairs showed low-to-moderate correlations, suggesting limited pairwise redundancy within the selected feature set.

**Table 3.**
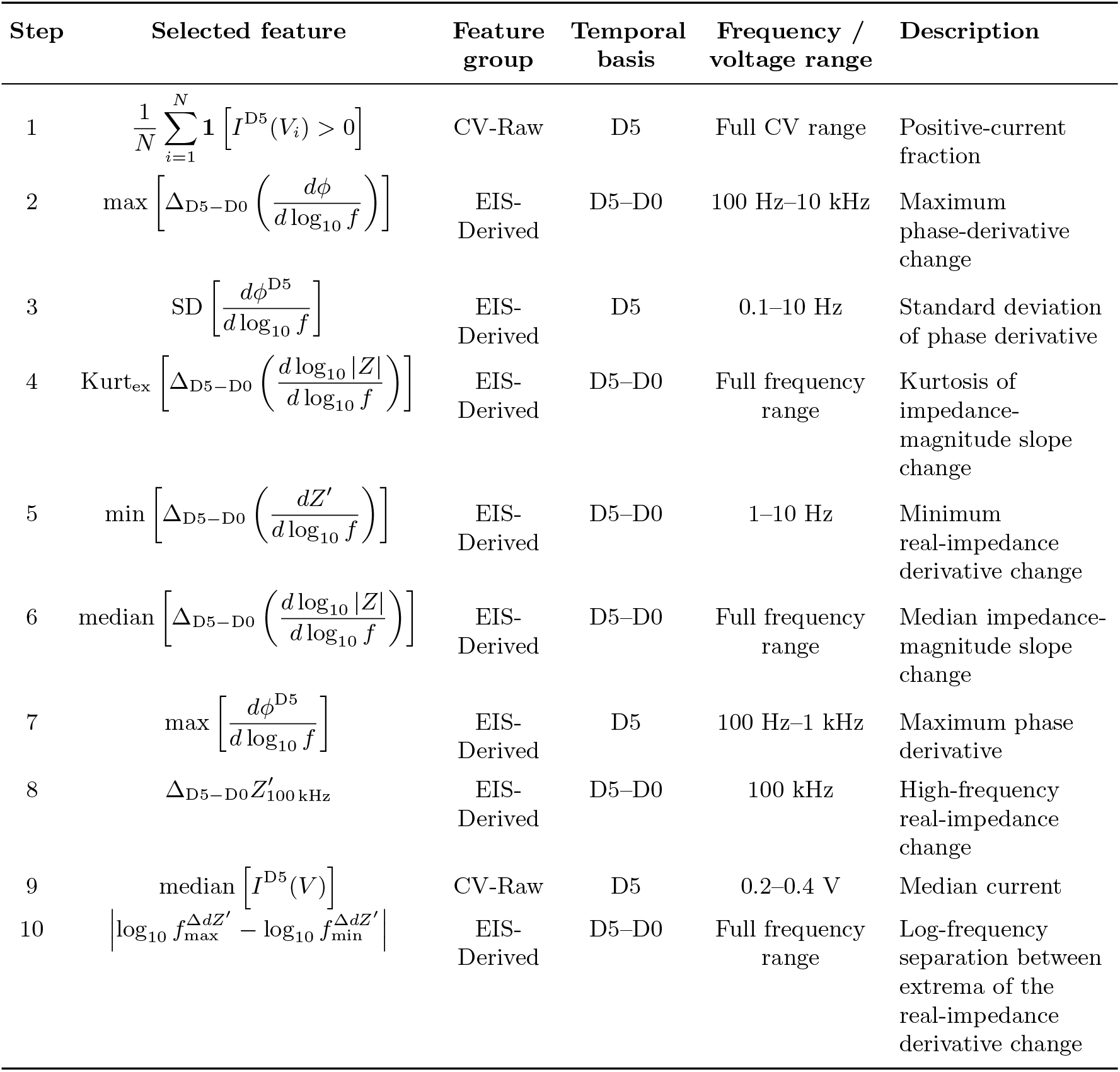
Selected early-life features obtained through stepwise forward feature selection.

**Figure 4.**
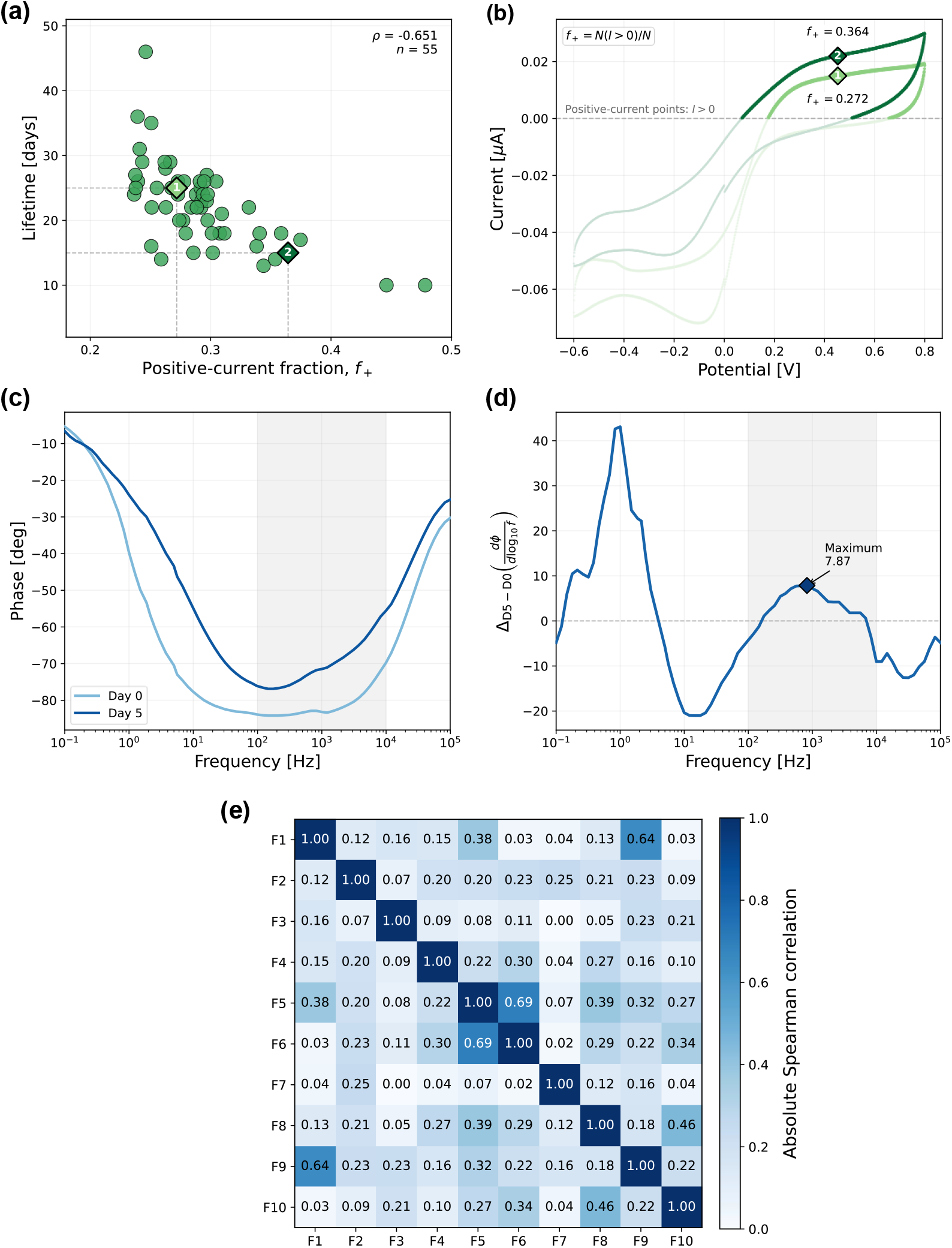
Early-life feature extraction and correlation analysis. (a) Relationship between the Day 5 positive-current fraction and lifetime for the 55 samples. T01M1 5 (1) and T01M2 2 (2) are highlighted for comparison. (b) Day 5 CV curves of the two highlighted channels, with positive-current portions emphasized to illustrate the positive-current fraction. (c) Day 0 and Day 5 EIS phase spectra of A9M3 4. The shaded region indicates the 100 Hz–10 kHz frequency range used for feature extraction. (d) Day 5 to Day 0 change in the phase derivative for A9M3 4. The diamond marks the maximum value within the 100 Hz–10 kHz range. (e) Absolute Spearman correlation matrix of the 10 selected early-life features. Feature labels F1–F10 correspond to Steps 1–10 in Table 3.

From an experimental perspective, the selected features may reflect changes in electrical continuity and electrode–electrolyte exposure during aging. Polymer–metal interfacial delamination can increase the electrochemically accessible area and alter the CV response, as reflected by increased CV area and enhanced hysteresis. [13] Such interfacial changes may also affect the shape and frequency dependence of the EIS spectra. Including Day 5 values and changes relative to Day 0 therefore allows the model to represent both the early electrochemical state and its evolution during aging. These descriptors provide information beyond the single 1 kHz impedance value used to define failure. However, the selected features alone cannot uniquely distinguish cracking, delamination, or other degradation mechanisms.

### 5.3 Overall Early-Lifetime Prediction Performance

After identifying the final 10 early-life features, we compared the predictive performance of different models using the same feature set. In addition to Elastic Net, we also evaluated Ridge, GPR, and LightGBM. No additional model-specific feature selection was performed.

Table 4 summarizes the overall prediction performance of the evaluated models. All machine learning models outperformed the dummy baseline, with Elastic Net achieving the lowest mean RMSE and MAPE of 3.711 accelerated aging days and 13.90%, respectively. Ridge achieved comparable performance, with an RMSE of 3.792 days and a MAPE of 14.19%. GPR and LightGBM showed higher prediction errors under the selected feature representation.

**Table 4.** Comparison of lifetime prediction performance across different models. Values represent mean ± standard deviation across five complete repetitions of five-fold device-level cross-validation.

| Model | $N_{\text{feature}}$ | RMSE [days] | MAPE (%) |
| --- | --- | --- | --- |
| Dummy baseline | 0 | $6.652 \pm 0.219$ | $25.85 \pm 1.13$ |
| Ridge | 10 | $3.792 \pm 0.343$ | $14.19 \pm 2.06$ |
| Elastic Net | 10 | $3.711 \pm 0.447$ | $13.90 \pm 2.22$ |
| GPR | 10 | $4.519 \pm 0.511$ | $17.79 \pm 2.73$ |
| LightGBM | 10 | $5.877 \pm 0.558$ | $22.72 \pm 1.75$ |

Given that the final feature subset was identified through an Elastic Net–based forward-selection procedure, this shared representation may favor Elastic Net. Its combination of coefficient shrinkage and variable selection provides a way to control model complexity when electrochemical predictors retain some correlation. [25] The similar performance of Ridge suggests that regularized linear models adequately represent the predictive relationships in this feature space. This does not imply that the underlying physical degradation processes are linear. Moreover, the development dataset contained 55 channels from 13 devices, so the observed model ranking should be interpreted within the present validation setting, recognizing the uncertainty associated with small-sample performance estimates. [26] Considering prediction performance, model complexity, and interpretability, Elastic Net was retained as the primary model for subsequent analyses.

Fig. 5 (a-c) presents the lifetime prediction results. Compared with LightGBM and GPR, Elastic Net predictions were generally closer to the identity line and showed a more concentrated residual distribution, consistent with Table 4. Nevertheless, some of the longest-lived channels remained underestimated. For reliability screening, this bias could undervalue particularly durable channels, emphasizing the importance of examining prediction errors across the lifetime range in addition to aggregate metrics. Fig. 5(d) presents the standardized coefficients of the 10 selected early-life features. The largest absolute coefficients corresponded to F5 (+2.36), F3 (+2.33), and F1 (−2.33). F5 describes the minimum change in the real-impedance derivative between Day 0 and Day 5 at 1 − 10 Hz, whereas F3 describes the variability of the Day 5 phase derivative at 0.1 − 10 Hz. These coefficients describe conditional associations within the fitted model. Their signs and magnitudes depend on the selected predictors and regularization, and should not be interpreted as direct measures of the contribution of individual physical failure mechanisms. [27]

**Figure 5.**
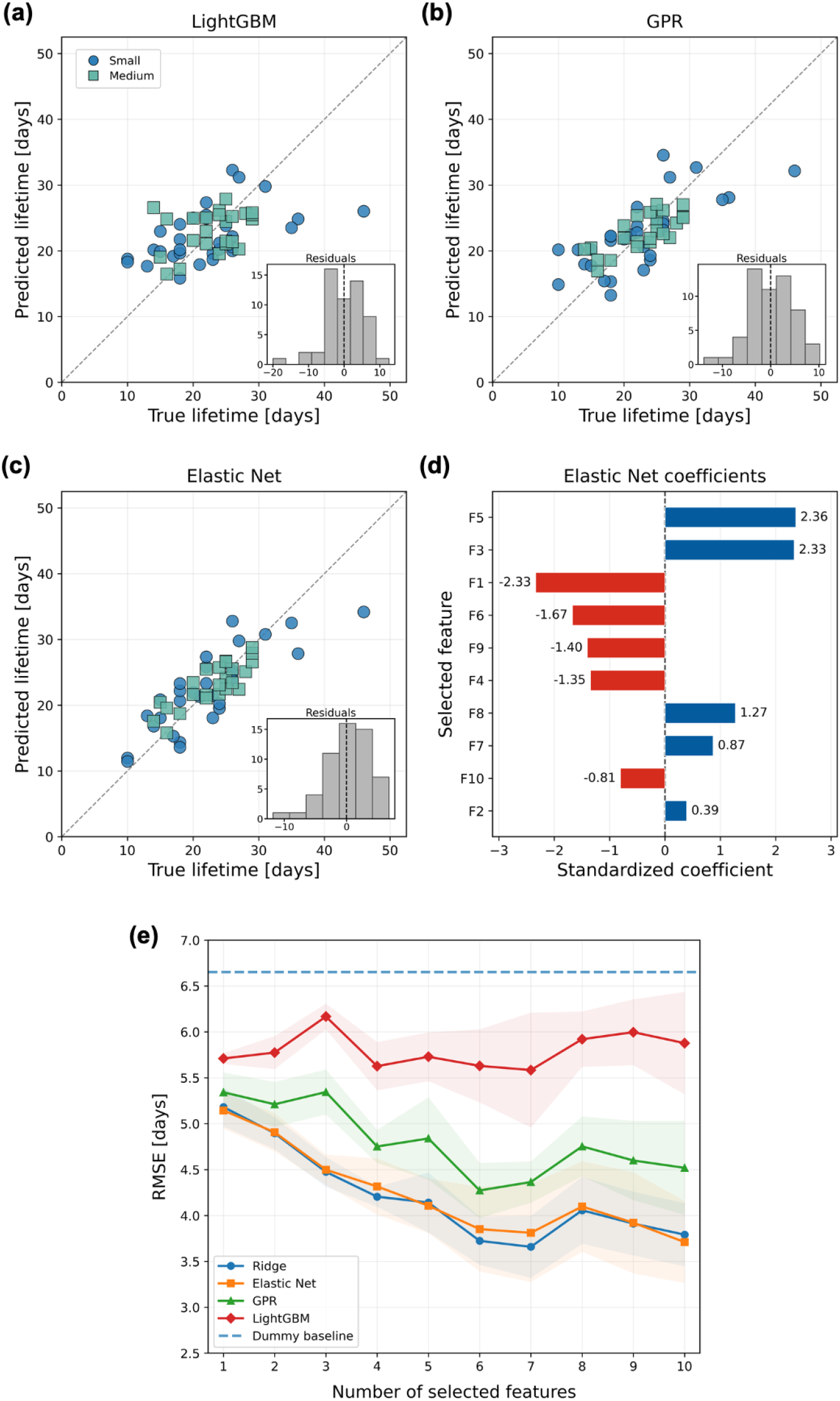
Overall early-lifetime prediction performance across models and feature coefficients of the Elastic Net model. (a–c) Parity plots for LightGBM, GPR, and Elastic Net, respectively, with samples grouped by electrode size. The dashed diagonals indicate perfect agreement between predicted and observed lifetimes. Insets show the residual distributions. (d) Standardized Elastic Net coefficients of the 10 selected early-life features, ranked by absolute magnitude. Positive and negative coefficients are shown in blue and red, respectively. Feature labels F1–F10 correspond to Steps 1–10 in Table 3. (e) Early-lifetime prediction performance as a function of feature count. The curves show RMSE, with shaded regions indicating ±1 standard deviation. Features were added sequentially according to a fixed forward-selection order, and the dashed line represents the dummy-regressor baseline. Lifetimes and RMSE are expressed in accelerated-aging days.

The prominence of these low-frequency descriptors provides an experimental rationale for examining spectral shape and its early evolution. Studies combining impedance analysis with electrode morphology have shown that changes in exposed conductive geometry and insulation condition influence the impedance response. [28] Longitudinal measurements have also demonstrated that spectral magnitude and phase can distinguish electrode conditions that are not adequately resolved by a single 1 kHz measurement. [29] Interpretation remains dependent on device configuration: electrode geometry, insulation thickness, and parasitic capacitance can influence measured impedance, particularly at higher frequencies. [30] Thus, the selected spectral features may represent several aspects of the device response rather than isolated resistance or capacitance parameters.

Both CV features, F1 and F9, received negative coefficients, associating higher feature values with shorter predicted lifetimes within the fitted model. F1 describes the positive-current fraction, while F9 describes the median current within 0.2–0.4 V. These features summarize selected aspects of the CV waveform rather than its integrated charge. Previous neural-electrode studies have reported CV broadening and increased charge-storage capacity consistent with developing leakage pathways. [31, 32] These findings provide a general rationale for examining CV-derived features in reliability prediction. However, the present analysis does not establish that F1 or F9 specifically represents CV broadening, charge-storage changes, or leakage. Their predictive associations may also reflect electrode geometry and initial electrochemical condition in the pooled Small and Medium dataset.

Fig. 5(e) shows prediction performance as features were added sequentially according to the selection order in Table 3. Most improvement for Elastic Net and Ridge occurred within the first 6–7 features, with smaller, nonmonotonic changes thereafter. Ridge reached its displayed minimum around seven features, whereas Elastic Net reached its minimum at ten. GPR also improved with additional features, while LightGBM benefited less.

Earlier neural-electrode studies established relationships between impedance measurements and subsequent recording functionality. [33] Here, the prediction target is the time to a defined impedance endpoint during accelerated aging. The results therefore support early reliability assessment under the studied conditions. Connecting these estimates to recording and stimulation performance remains necessary, particularly because human explant studies have demonstrated material-dependent relationships between impedance, physical damage, and electrode function. [34]

### 5.4 Performance Analysis across Device Characteristics

To further evaluate prediction performance across different device characteristics, we analyzed Elastic Net errors at the device level. Fig. 6 shows the RMSE for each device relative to the dummy baseline. Devices are grouped by electrode size (Small and Medium) and coating condition (Control, ALD, and ALI). Detailed predictions for individual devices and channels are provided in the Supplementary Material.

**Figure 6.**
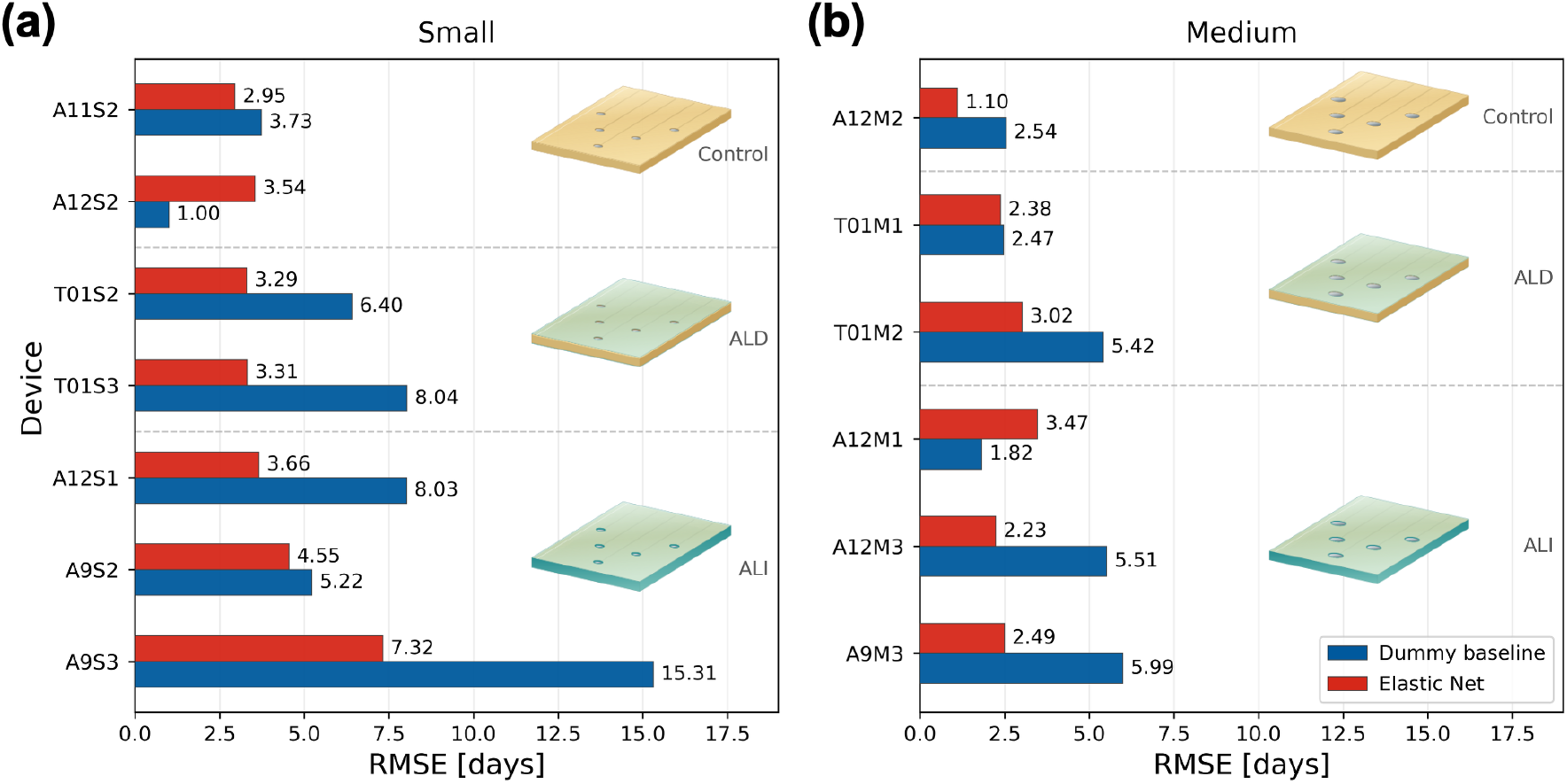
Device-level prediction errors across electrode sizes and coating conditions. RMSE, expressed in accelerated aging days, is shown for the Elastic Net model (red) and the dummy baseline (blue) for (a) Small electrodes and (b) Medium electrodes, with devices grouped by coating condition (Control, ALD, and ALI). Insets schematically illustrate the MEA electrode regions: amber represents polyimide (PI), gray represents exposed platinum sites, and green indicates inorganic barrier coverage. Control devices retain PI insulation without an additional inorganic barrier. ALD covers the top and bottom faces, whereas ALI-based encapsulation additionally protects the outer edges and electrode-via sidewalls. Schematics are not to scale.

Overall, Elastic Net achieved lower RMSE than the dummy baseline for 11 of the 13 devices, with A12S2 and A12M1 as the exceptions. For these devices, channel lifetimes were tightly clustered around the dummy baseline’s constant prediction; for example, A12S2 lifetimes ranged only from 22 to 24 accelerated aging days. This lifetime distribution favored the mean-based dummy baseline, and Elastic Net did not further reduce the prediction error. Performance relative to this baseline therefore depends partly on each device’s lifetime distribution.

Small electrodes generally showed higher and more variable prediction errors, suggesting that lifetime prediction was more challenging for this group. Differences across coating conditions were most evident among Small electrodes, where ALI devices showed higher errors than Control and ALD devices. Within the Medium group, however, ALI and ALD errors overlapped. These descriptive differences should be interpreted cautiously because each size–coating subgroup contained only one to three devices.

The encapsulation architectures provide useful experimental context for these comparisons. Control devices retained their polyimide insulation without an additional inorganic barrier. Conventional ALD devices incorporated alumina/titania barriers on the top and bottom surfaces, whereas the 3D-ALI approach combined alumina infiltration with a titania cap applied after device release, extending protection to polymer sidewalls. The earlier experimental study also demonstrated greater bending tolerance for ALI coatings under the conditions tested, although these coating-level improvements did not establish a general extension of device lifetime. Metal cracking and interlayer delamination remained important competing failure modes. [11]

Adding inorganic or organic/inorganic composite barriers introduces mechanical considerations alongside permeability benefits. [35, 36, 37] Crack onset strain depends on film thickness and loading mode. [38, 39] Studies of related inorganic silicon nitride encapsulation/polymer substrate systems have demonstrated moisture-assisted crack growth at strains below the measured crack onset strain. [40] Damage in the underlying polymer can also alter the subsequent evolution of barrier cracking. [41] These findings illustrate why short-term strain tolerance alone cannot fully describe long-term mechanical reliability. Related work on PECVD amorphous silicon carbide has shown that deposition conditions influence oxidation susceptibility and the evolution of residual stress during environmental exposure. [42]

Fabrication-dependent adhesion is another relevant factor. In polyimide-based neural-implant test structures, changing the curing conditions of the underlying polyimide layer substantially altered insulation lifetime in saline. [43] Separate adhesion measurements showed that the type of surface treatment strongly influenced bonding between polyimide layers. [44] Susceptibility to delamination therefore depends on the specific interfaces and processing history as well as the barrier material. The ALD and ALI groups examined here represent particular combinations of materials and fabrication steps, rather than an isolated comparison of deposition and infiltration processes.

Fabrication can also affect the early electrochemical inputs before substantial aging occurs. The earlier study attributed higher initial impedance in ALI electrodes partly to a residual oxide ring around the electrode perimeter. [11] Such differences in effective exposed area and initial electrochemical state may influence the engineered features, but their contribution to the observed prediction errors was not isolated. Higher RMSE should therefore not be interpreted as evidence of poorer encapsulation reliability. Establishing how processing, mechanical damage, and interfacial degradation individually affect prediction performance requires additional characterization beyond the scope of this study.

### 5.5 Cross-Batch Long-Lifetime Prediction under Distribution Shift

Compared with Batch 1, Batch 2 showed generally longer lifetimes. The increase was particularly pronounced for the Medium group, while the Small group also shifted toward a longer and more concentrated lifetime range. This comparison therefore evaluates whether relationships learned from the development batch remain useful for predicting the longer lifetimes observed in independently fabricated devices. To examine whether this difference was also reflected in the early-life feature space, t-SNE [45] visualization was performed using the final 10 selected features. As shown in Fig. 7(a), the Batch 2 Small and Medium samples were separated from the Batch 1 data, consistent with differences in their early electrochemical feature distributions.

**Figure 7.**
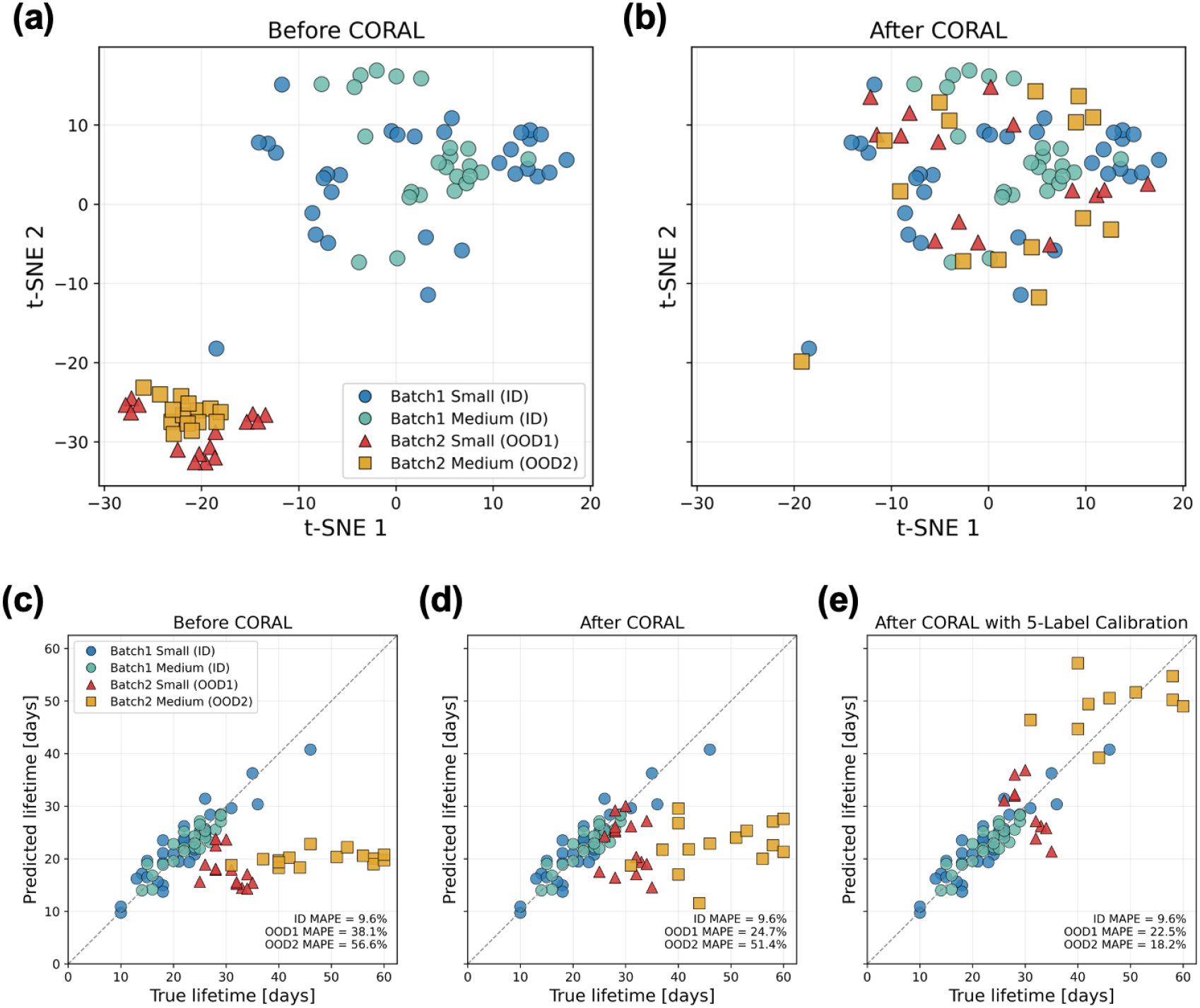
Cross-batch feature alignment and lifetime prediction using CORAL and limited-label calibration. (a,b) t-SNE visualizations of the 10 selected early-life features before and after CORAL alignment, respectively. (c–e) Predicted versus observed lifetimes using (c) the source-domain Elastic Net model directly, (d) CORAL alignment, and (e) CORAL alignment followed by calibration using five labeled channels from one device in each Batch 2 electrode-size group. Each point represents one electrode channel. ID denotes the Batch 1 development data, while OOD1 and OOD2 denote the Batch 2 Small and Medium groups, respectively. Dashed diagonal lines in (c–e) indicate perfect agreement between predicted and observed lifetimes. Lifetimes are expressed in accelerated aging days.

Because fabrication and testing conditions both varied between batches, the observed differences cannot be attributed to a single experimental factor. The possible contributions of handling and failure behavior are discussed in Section 4.2. In addition, the EIS frequency-sampling density differed between batches, as described in Section 2.1. This acquisition difference may influence spectral derivatives and other engineered descriptors, emphasizing the importance of consistent measurement and preprocessing protocols when transferring models between datasets.

To reduce the impact of the observed distribution shift, an unsupervised adaptation approach was first applied without using target lifetime labels. Specifically, CORAL (CORrelation ALignment) was used to reduce differences between the covariance structures of the source and target features. [46] After alignment, the Batch 2 samples showed greater overlap with the Batch 1 data in the t-SNE visualization in Fig. 7(b). However, this visual overlap alone does not establish that the source-domain model accurately predicts target-domain lifetimes.

For cross-batch evaluation, the Elastic Net model was trained on the full Batch 1 development set (55 channels), achieving a training MAPE of 9.6% on the source data. Direct application of the source-domain model resulted in MAPE values of 38.1% for OOD1 and 56.6% for OOD2, with a general tendency to underestimate longer lifetimes, as shown in Fig. 7(c). After CORAL alignment, MAPE decreased to 24.7% for OOD1 but only slightly to 51.4% for OOD2 (Fig. 7(d)). Thus, covariance alignment improved transfer for the Small group, while substantial underprediction remained for the longer-lived Medium group. Matching the input covariance structure alone was insufficient to remove this prediction bias.

Calibration was subsequently evaluated using five labeled channels from one of the three devices in each target group. After CORAL alignment and calibration, the reported MAPE values decreased to 22.5% for OOD1 and 18.2% for OOD2 (Fig. 7(e)). The particularly large improvement for the Medium group indicates that target lifetime information helped correct prediction bias that unsupervised alignment alone did not resolve. This result supports combining feature alignment with limited target-domain calibration when transferring predictions to a new fabrication batch.

The experimental requirements of calibration should nevertheless be recognized. Obtaining lifetime labels requires observing the reference channels until they reach the defined failure endpoint, so calibration retains a need for complete aging tests on reference devices. Furthermore, the five channels in each group belong to a single device and share fabrication and aging conditions. Additional independent reference devices and batches are therefore needed to establish how reliably this calibration strategy transfers beyond the present datasets.

### 5.6 Discussion and limitations

The practical value of early-life prediction lies in obtaining useful reliability information before every device has reached failure. Following broader validation for a defined device family and testing protocol, this approach could support earlier selection of fabrication conditions or design candidates for extended reliability studies. Its applicability to other implantable microelectronic devices would depend on identifying accessible early measurements that respond to the relevant degradation processes. Depending on the device, these measurements could include insulation leakage current or interconnect resistance, with prediction targets defined by the electrical requirements of the intended application. Selecting such indicators should be guided by the expected failure mechanisms and service loading, consistent with physics-of-failure approaches to electronic prognostics. [47]

The statistical scope of the present study remains limited. Model development used 55 channels from 13 devices, and cross-batch evaluation involved only one additional fabrication batch. Device-level partitioning reduces information leakage between related channels, but the number of independently fabricated devices remains small. Large electrodes and four channels with premature bond-pad cracking were excluded from model development, limiting conclusions about those geometries and early failures. Prospective validation on additional batches should assess prediction uncertainty and the stability of the selected features. Studies targeting substantially longer lifetimes should also accommodate channels that remain operational when testing ends through methods that account for right-censored lifetime data. [48]

The prediction target is the time to a specified impedance change during accelerated aging, which does not directly establish functional lifetime in vivo. Testing at a single aging temperature of 87°C does not establish an acceleration relationship with physiological conditions. Studies of several polyimide formulations have reported changes in mechanical properties during PBS exposure at 85°C, despite comparative stability at 37°C and 60°C. [49] Validation at multiple temperatures, including physiological temperature, is needed to determine whether the same early signatures and failure processes remain relevant. The surrounding chemical environment also requires consideration, as reactive accelerated-aging studies have demonstrated distinct effects of temperature and oxidative exposure on implant degradation. [18] Connecting predicted lifetimes to recording quality, stimulation performance, and viable-channel fraction would help establish their relevance to neural-interface function.

A further priority is to define the combinations of device architecture, processing, and loading conditions under which early prediction remains reliable. Controlled studies could vary barrier encapsulation thickness, infiltration conditions, curing, and surface preparation while measuring strain history, crack development, and interfacial adhesion. Combining these measurements with electrochemical monitoring and microscopy could distinguish whether selected features precede physical damage, track its progression, or primarily reflect differences in the initial device state. This would enable failure-mode-specific interpretation without treating ALD or ALI as a general indicator of reliability. Such characterization is beyond the scope of the present prediction study, but would help define its range of applicability and identify conditions requiring recalibration or additional measurements.

## 6 Conclusion

This study demonstrates early prediction of neural microelectrode lifetime under accelerated aging using EIS and CV features combined with Elastic Net regression. Using measurements collected through Day 5, the model achieved an RMSE of 3.71 accelerated-aging days and a MAPE of 13.90%, while covariance alignment and calibration with limited labeled data improved predictions across fabrication batches. The findings are limited by the small dataset, a single accelerated-aging temperature, and an impedance-based failure criterion that does not directly establish in vivo functional lifetime. Future work should validate the approach across larger independent datasets, device architectures, and aging conditions, quantify prediction uncertainty, and relate predicted lifetimes to recording and stimulation performance.

## Acknowledgments

Hui Hua, Tingkai Li, and Chao Hu acknowledge partial research support from a MathWorks Development Collaborative Research Grant (DCRG). Kyungjin Kim acknowledges partial research support from Blackrock Neurotech (Blackrock Microsystems, Inc.) and the National Research Foundation of Korea (NRF) (Grant No. RS-2024-00347831). Chao Hu and Kyungjin Kim acknowledge partial research support from the U.S. Army DEVCOM Ground Vehicle Systems Center (GVSC) under Cooperative Agreement No. W56HZV-21-2-0001 through the VIPR-GS program.

## Author contributions

Conceptualization: K.K. and C.H. Methodology: H.H., T.L., C.H., and K.K. Software and formal analysis: H.H. and T.L. Investigation: M.N., R.M., and H.J. Data curation: H.H., M.N., R.M., and H.J. Visualization: H.H., K.K., and C.H. Supervision and funding acquisition: C.H. and K.K. Writing - original draft: H.H. Writing - review and editing: H.H., M.N., T.L., R.M., H.J., K.K., and C.H. All authors have read the manuscript and provided comments.

## Data availability

The data supporting the findings of this study are available in the article, its supplementary material, and the Zenodo repository at https://doi.org/10.5281/zenodo.23679832. This study used ModularML, an open-source machine learning framework, to support model development [50].

## Supplementary data

Supplementary material for this article is available online.

