## Supporting Information for "Machine Learning-Based Early Prediction of Neural Microelectrode Lifetime Under Accelerated Aging Conditions"

This supplement describes the experimental aging dataset and supporting methods used for early prediction of microelectrode lifetime.

**Contents**

S1 Dataset description

S2 Detailed early-life feature engineering

S3 Machine learning settings and validation

References

### S1 Dataset description

#### S1.1 Dataset origin and composition

The experimental dataset combines electrochemical aging measurements previously reported by Niemiec et al. [1] with measurements from additional devices fabricated for the present study. The published study tested 17 polyimide microelectrode arrays: four uncoated controls, five arrays with conventional atomic layer deposition (ALD) encapsulation, and eight arrays with three-dimensional atomic layer infiltration (3D-ALI) encapsulation. Each array contained five platinum microelectrode sites. Electrode geometric surface areas were 1,000, 10,000, or 100,000 $\mu m^2$, referred to as Small, Medium, and Large, respectively. Table 2 of the published study reports 77 channel records for the impedance-defined failure endpoint: 31 Small, 28 Medium, and 18 Large channels. These counts refer specifically to the impedance-endpoint dataset; the published study also analyzed a separate endpoint based on combined electrochemical and visual evidence of total failure. Some sites did not meet the impedance endpoint before testing was stopped. The present manuscript describes 15 additional channels of each electrode size, giving the collection summarized in Table S1.

**Table S1. Experimental channel counts described in Section 2.1 of the main manuscript.**

| Electrode size | Original channels | Additional channels | Combined channels |
|---|---|---|---|
| Small, 1,000 $\mu m^2$ | 31 | 15 | 46 |
| Medium, 10,000 $\mu m^2$ | 28 | 15 | 43 |
| Large, 100,000 $\mu m^2$ | 18 | 15 | 33 |
| Total | 77 | 45 | 122 |

The primary prediction analysis uses Small and Medium channels. As specified in Table 2 of the main manuscript, the Batch 1 development set contains 55 channels from 13 devices. The independently fabricated Batch 2 test sets contain 15 Small channels from three devices and 15 Medium channels from three devices, giving 85 modeled channels from 19 devices in total. Large electrodes are outside the primary model-development set because the published study found greater susceptibility to cracking at the electrode-trace junction.

#### S1.2 Device structure and fabrication

In the published cohort, the electrode sites and conductive traces were embedded between base and top layers of PI2610 polyimide, each reported as approximately 2-3 μm thick. The metal stack was deposited by electron-beam evaporation and consisted of Cr/Pt/Au/Pt/Cr with nominal thicknesses of 30/20/100/20/20 nm. Chromium promoted adhesion, while the platinum layers separated gold and chromium during subsequent thermal processing. The upper chromium layer was removed locally at the electrode openings to expose platinum as the active electrode surface. The base polyimide was treated with oxygen reactive ion etching before metal patterning to improve interfacial adhesion. After the top polyimide layer was deposited and cured, electrode openings and device outlines were patterned using photolithography and plasma etching. Wires were attached to the bond pads, the contact region was sealed with silicone, and devices were released by dissolving the sacrificial magnesium layer. Detailed fabrication recipes and the full process sequence are provided in Sections 2.1-2.3 and Figure 2 of the published paper [1].

#### S1.3 Encapsulation groups

The Control group consisted of polyimide arrays without an added metal-oxide barrier. For conventional ALD devices, alumina/titania coatings were incorporated on the top and bottom while the devices remained on the wafer. Subsequent etching of electrode openings and device outlines left exposed polymer sidewalls. For 3D-ALI devices, released arrays were suspended vertically in the reactor, allowing precursor access to both faces, device edges, and electrode-opening sidewalls. Alumina infiltration created a polymer-inorganic hybrid region, followed by a titania cap. Sacrificial photoresist over the electrode sites was subsequently removed to expose active platinum while retaining the surrounding encapsulation. All ALD/ALI processes in the published study were performed at 120 °C. Alumina ALI used trimethylaluminum (TMA) and water, a target pressure of 6 Torr, a 3 min diffusion interval for each precursor, and a 6 min purge interval, repeated for 60 cycles. The titania cap was deposited using tetrakis(dimethylamino)titanium (TDMAT) and water; TDMAT was heated to 75 °C, and 525 ALD cycles targeted an approximately 30 nm titania film. The conventional on-wafer coating sequence used 60-cycle alumina and 525-cycle titania layers [1].

#### S1.4 Accelerated aging and electrochemical measurements

Devices were maintained in phosphate-buffered saline (PBS) at 87 °C between measurements. Electrochemical measurements were performed in PBS maintained at 37 °C in a heated water bath. A three-electrode configuration used the microelectrode site as the working electrode, an Ag/AgCl reference electrode, and a platinum-wire counter electrode. Measurements were acquired using a CHI 660E electrochemical workstation (CH Instruments). EIS spectra were collected from 0.1 Hz to 100 kHz with a 50 mV excitation amplitude. CV measurements swept the potential between -0.6 and +0.8 V versus Ag/AgCl at 100 mV/s. Three consecutive CV cycles were recorded in the published protocol, and the final cycle was used for analysis. Daily impedance magnitude and phase at 1 kHz, charge per phase derived from CV, and optical images supported longitudinal assessment of degradation [1].

For optical imaging, devices were briefly rinsed with deionized water after electrochemical testing, dried, and held flat for imaging before being returned to their PBS vials and the 87 °C oven. The current manuscript reports less frequent imaging in Batch 2 and reduced EIS sampling density for the additional devices: 12 points per frequency decade in the original dataset, six points per decade during the first week for the new devices, and three points per decade thereafter. These changes reduced measurement time while retaining information for feature extraction and failure assessment. Possible effects of handling differences on cracking remain qualitative and were not independently isolated.

#### S1.5 Failure endpoint and lifetime labels

The impedance-based failure criterion and lifetime definition are provided in Section 2.2 of the main manuscript. Lifetimes are expressed as elapsed days of accelerated aging at 87 °C, without conversion to in vivo functional lifetime.

#### S1.6 Data files and channel inventory

The EIS and CV data are available in the repository identified in the Data availability statement of the main manuscript. The accompanying README describes the file organization, variables, units, and device/channel identifiers.

### S2 Detailed early-life feature engineering

#### S2.1 Electrochemical data preprocessing

EIS and CV measurements from Day 0 and Day 5 were used for feature extraction. For each microelectrode channel, Day 5 measurements were used to construct fixed-day features, while Day 0 measurements served as the baseline for cross-day features. No measurements collected after Day 5 were used.

For EIS data, frequency, real and imaginary impedance, impedance magnitude, and phase were obtained from the original measurement files. Invalid data points and those with nonpositive frequencies or impedance magnitudes were removed. For raw EIS features, impedance magnitude and phase were linearly interpolated along the logarithmic frequency axis to 10 predefined frequencies ranging from 0.1 Hz to 100 kHz. In contrast, derived EIS features were extracted directly from the cleaned full spectra.

For CV data, potential and current values were extracted, and invalid data points were removed. Up to the last 2,800 valid data points from each measurement were used for feature extraction. Raw CV features were calculated directly from these measurements. For derived CV features, the current signal was smoothed using a Savitzky–Golay filter with a window length of 31 points and a polynomial order of 3 before derivative calculation. Further details are provided in Section S2.3.2.

#### S2.2 EIS-based feature extraction

EIS-based features were divided into raw and derived features. Raw features were extracted from the impedance magnitude and phase spectra, while derived features were constructed from spectral derivatives and Nyquist representations.

##### S2.2.1 Raw EIS features

Raw EIS features were extracted from the impedance magnitude and phase spectra interpolated at 10 predefined frequencies, as described in Section S2.1. These included fixed-frequency values, global statistical descriptors, integral-based descriptors, and frequency-band features. The feature definitions, calculation methods, and corresponding feature counts are summarized in Table S2.

##### S2.2.2 Derived EIS features

Derived EIS features were extracted from the full EIS spectra using derivative-based and Nyquist-based descriptors. Five derivative curves were calculated numerically with respect to logarithmic frequency, and statistical, extreme-value, and frequency-band features were extracted from each curve. The derivative definitions and extracted descriptors are summarized in Table S3.

Additional features were extracted from the Nyquist representation to characterize impedance geometry and frequency-dependent behavior. These included geometric anchors, peak descriptors, impedance spans, arc characteristics, and low-frequency tail features. Detailed definitions and calculations are provided in Table S4.

**Table S2. Summary of raw EIS feature definitions.**

| Feature category | Descriptors | Calculation | Number |
|---|---|---|---|
| Fixed-frequency values | Impedance magnitude, phase | Values at 10 predefined frequencies | 20 |
| Global statistics | Mean, SD, variance, min, max, median, range, skewness, excess kurtosis | Calculated over the interpolated spectrum | 18 |
| Global integral | Area | Trapezoidal integration over $\log_{10}(f)$ | 2 |
| Frequency-band statistics | Mean, SD, min, max | Calculated within each frequency band | 80 |
| Frequency-band shape | Slope, area | Linear fitting and trapezoidal integration | 40 |
| Total | - | - | 160 |

*Frequency bands: 0.1–1, 1–10, 10–100, 100–1,000, 1,000–10,000, 10,000–100,000, 0.1–10, 1–100, 10–1,000, and 100–10,000 Hz.
*Predefined frequencies: 0.1, 0.3, 1, 3, 10, 30, 100, 1,000, 10,000, and 100,000 Hz.
*Slope calculation: For impedance magnitude, the slope was obtained by linear regression of $\log_{10}|Z|$ against $\log_{10}(f)$; for phase, the slope was obtained by regression of $\phi$ against $\log_{10}(f)$.

**Table S3. Definitions of EIS derivative features.**

(a) Derivative definitions

| Derivative feature | Mathematical definition |
|---|---|
| Real impedance derivative | $\frac{dZ'}{d\log_{10}(f)}$ |
| Imaginary impedance derivative | $\frac{dZ''}{d\log_{10}(f)}$ |
| Impedance magnitude derivative | $\frac{d\|Z\|}{d\log_{10}(f)}$ |
| Phase derivative | $\frac{d\phi}{d\log_{10}(f)}$ |
| Log-magnitude derivative | $\frac{d\log_{10}\|Z\|}{d\log_{10}(f)}$ |

(b) Extracted descriptors

| Feature category | Descriptors | Number |
|---|---|---|
| Global statistics and integral | Mean, SD, variance, min, max, median, range, skewness, excess kurtosis, signed area | 10 |
| Additional extreme-value descriptors | Frequencies at max/min, peak-to-valley amplitude, log-frequency separation, absolute area | 5 |
| Frequency-band descriptors | Mean, SD, min, max, slope, area across 10 bands | 60 |
| Total | - | 75 |

**Table S4. Definitions of Nyquist-based EIS features.**

| Feature category | Descriptors | Number |
|---|---|---|
| Geometric anchors | $Z'$ at highest and lowest frequencies; $-Z''$ at lowest frequency | 3 |
| Peak descriptors | Maximum $-Z''$, corresponding $Z'$and frequency | 3 |
| Impedance spans | Range of $Z'$ and $-Z''$ | 2 |
| Nyquist integral | $\int(-Z'')\,dZ'$ | 1 |
| Arc descriptors | Height: $\max(-Z'')$; Width: $\mathrm{W} = \max(Z') - \min(Z')$; Aspect ratio: Height/Width | 3 |
| Low-frequency tail | Tail slope: linear regression of $-Z''$ against $Z'$ using the five lowest-frequency points; Tail ratio: $-Z''/Z'$ at the lowest frequency | 2 |
| Global statistics | Mean, SD, variance, min, max, median, range, skewness, excess kurtosis, and area for $Z'$ and $-Z''$ | 19 |
| Frequency-band descriptors | Mean, SD, min, max, slope, and area across 10 frequency bands for $Z'$ and $-Z''$ | 120 |
| Total | - | 153 |

*The Nyquist integral was calculated using signed trapezoidal integration, with data points ordered by ascending frequency. The global statistics count excludes the maximum $-Z''$, which is already included among the peak descriptors.

#### S2.3 CV-based feature extraction

CV-based features were divided into raw and derived features. Raw features were extracted from the current–potential curves, while derived features were calculated from the differential current response ($dI/dV$).

##### S2.3.1 Raw CV features

Raw CV features were extracted from the current–potential measurements described in Section S2.1. These included global statistical descriptors, integral-based descriptors, current polarity features, extreme-value features, and voltage-window descriptors. The feature definitions, calculation methods, and corresponding feature counts are summarized in Table S5.

##### S2.3.2 Derived CV features

Derived CV features were extracted from the differential current response ($dI/dV$). Following Savitzky–Golay smoothing, $dI/dV$ was calculated as the ratio of the numerical current gradient to the numerical potential gradient. Data points with absolute potential gradients of $10^{-6}$ or less were excluded to avoid unstable derivative values.

**Table S5. Summary of raw CV feature definitions.**

| Feature category | Descriptors | Number |
|---|---|---|
| Global statistics | Mean, SD, variance, min, max, median, range, skewness, excess kurtosis | 9 |
| Global integrals and polarity | Signed area, absolute area, positive and negative current areas, positive and negative current fractions | 6 |
| Additional extreme-value descriptors | Potentials at max/min current, peak-to-valley amplitude, potential separation between extrema | 4 |
| Voltage-window statistics | Mean, SD, min, max, median, range across 12 windows | 72 |
| Voltage-window shape descriptors | Slope, signed area, absolute area across 12 windows | 36 |
| Total | - | 127 |

*The 12 predefined potential windows were −0.6 to −0.4, −0.4 to −0.2, −0.2 to 0, 0 to 0.1, 0.1 to 0.2, 0.2 to 0.4, 0.4 to 0.6, 0.6 to 0.8, −0.6 to −0.2, −0.2 to 0.2, 0 to 0.4, and 0.2 to 0.8 V. Areas were calculated by trapezoidal integration after sorting the data by potential. Slopes were obtained by linear regression of current against potential.

From each $dI/dV$ curve, global statistical descriptors, integral-based descriptors, extreme-value features, and voltage-window descriptors were extracted. Their definitions and corresponding feature counts are summarized in Table S6.

**Table S6. Summary of derived CV feature definitions.**

| Feature category | Descriptors | Number |
|---|---|---|
| Global statistics | Mean, SD, variance, min, max, median, range, skewness, excess kurtosis | 9 |
| Global integrals | Signed area, absolute area | 2 |
| Additional extreme-value descriptors | Potentials at max/min $dI/dV$, peak-to-valley amplitude, potential separation between extrema | 4 |
| Voltage-window statistics | Mean, SD, min, max, median, range across 15 windows | 90 |
| Voltage-window shape descriptors | Slope, signed area, absolute area across 15 windows | 45 |
| Total | - | 150 |

*The 15 predefined potential windows included the 12 windows listed in Table S5 and three additional windows: −0.1 to 0.1, 0 to 0.2, and 0.1 to 0.3 V. Window-based descriptors were calculated from the valid $dI/dV$ values within each potential range. Areas were computed by trapezoidal integration with respect to potential, and slopes were obtained by linear regression of $dI/dV$ against potential.

#### S2.4 Cross-day feature construction

In addition to fixed-day features, cross-day features were constructed to characterize changes in electrochemical responses between Day 0 and Day 5. These features were intended to capture early aging behavior that may not be fully reflected in measurements from a single time point.

For raw EIS features, differences and ratios of impedance magnitude and differences in phase were first calculated, followed by the extraction of statistical and frequency-band descriptors. Derived EIS features were constructed from changes in spectral derivatives and Nyquist-based descriptors. For CV data, cross-day features were calculated as differences between corresponding features extracted from Day 5 and Day 0. The calculation methods and feature counts are summarized in Table S7.

**Table S7. Summary of cross-day feature construction.**

| Feature group | Feature category | Calculation | Number |
|---|---|---|---|
| Raw EIS | Impedance magnitude difference | $\Delta\lvert Z\rvert = \lvert Z_5\rvert - \lvert Z_0\rvert$ | 80 |
| | Impedance magnitude ratio | $R_{\lvert Z\rvert} = \lvert Z_5\rvert / \lvert Z_0\rvert$ | 80 |
| | Phase difference | $\Delta\phi = \phi_5 - \phi_0$ | 80 |
| Derived EIS | Spectral derivative changes | $\Delta D_q = D_{q,5} - D_{q,0}$ | 375 |
| | Nyquist component changes | $\Delta Z' = Z'_5 - Z'_0$<br>$\Delta(-Z'') = (-Z''_5) - (-Z''_0)$ | 140 |
| | Nyquist geometric changes | $\Delta g = g_5 - g_0$<br>$R_g = g_5/g_0$ | 28 |
| Raw CV | Feature differences | $\Delta h_{\text{raw}} = h_{\text{raw},5} - h_{\text{raw},0}$ | 127 |
| Derived CV | Feature differences | $\Delta h_{\text{derived}} = h_{\text{derived},5} - h_{\text{derived},0}$ | 150 |
| Total | - | - | 1060 |

*$D_q$, $g$, and $h$ represent spectral derivatives, Nyquist geometric descriptors, and CV features, respectively. For EIS curve-based features, changes were calculated before feature extraction, whereas Nyquist geometric and CV features were extracted before calculating changes.

### S3 Machine learning implementation and evaluation

The main text presents the proposed machine learning framework for early prediction of neural microelectrode lifetime and summarizes its predictive performance. This study used ModularML, an open-source machine learning framework, to support model development [2]. This section provides additional implementation and evaluation details.

#### S3.1 Feature Screening and Selection

Additional implementation details for the feature screening and forward feature selection procedures described in Section 4.3 of the main text are summarized in Table S8.

**Table S8. Implementation settings for forward feature selection.**

| Component | Parameter | Setting / Search space |
|---|---|---|
| Preprocessing | Standardization | StandardScaler |
| Cross-validation | Strategy | 5-fold device-level CV |
| | Random seed | 2026 |
| Selection model | Algorithm | Elastic Net |
| Hyperparameter tuning | $\lambda$ | $10^{-4}$ - $10^{2}$ |
| | $\alpha$ | 0.1, 0.3, 0.5, 0.7, 1.0 |
| Feature selection | Maximum features | 10 |
| | Selection criterion | Minimum mean CV RMSE |
| | Feature count criterion | Minimum (mean CV RMSE + SD) |

**Table S9. Model configurations and hyperparameter search spaces.**

| Model | Hyperparameter | Search space / Setting |
|---|---|---|
| Ridge | $\lambda$ | $10^{-4}$ - $10^{3}$ |
| Elastic Net | $\lambda$ | $10^{-4}$ - 1 |
| | $\alpha$ | 0.1, 0.3, 0.5, 0.7, 0.9 |
| GPR | Kernel | RBF, Matérn ($\upsilon$=1.5, 2.5) |
| | Noise regularization | $10^{-8}$, $10^{-6}$, $10^{-4}$ |
| LightGBM | Number of estimators | 50, 100 |
| | Learning rate | 0.03, 0.10 |
| | Number of leaves | 3, 7 |
| | Minimum child samples | 3, 5 |

#### S3.2 Model Training and In-Distribution Evaluation

Forward feature selection was performed once on the full Batch 1 development dataset using a single five-fold device-level cross-validation. The resulting 10-feature subset was held fixed across all outer cross-validation folds and all five repetitions for the subsequent model comparisons. All models were evaluated using the same 10 selected features and repeated five-fold device-level cross-validation (five repetitions). Within each outer training fold, hyperparameters were optimized using an inner five-fold device-level cross-validation. Prediction performance was calculated from the out-of-fold predictions covering all 55 channels in each repetition. Table 4 of the main manuscript reports the mean ± standard deviation of these performance metrics across the five complete repetitions.

#### S3.3 Cross-Batch Evaluation and Domain Adaptation

For cross-batch evaluation, the Elastic Net model was trained on all 55 Batch 1 channels using the 10 selected features, with hyperparameters optimized through five-fold device-level cross-validation. The trained model was applied directly to the Batch 2 Small and Medium groups for source-only prediction. CORAL was then applied separately to each target group, aligning its feature distribution with the full Batch 1 reference set using unlabeled target data.

For calibration, five labeled channels from one device in each target group were used to calculate a size-specific correction based on the median prediction residual:

$$b_s = \text{median}_{j \epsilon C_s}\left(L_j - \hat{L}_j^{\text{CORAL}}\right) \tag{1}$$

where $C_s$ denotes the five calibration channels in group $s$, and $L_j$ and $\hat{L}_j^{\text{CORAL}}$ are the observed and CORAL-based predicted lifetimes, respectively. The calibrated predictions were then obtained as

$$\hat{L}_i^{\text{cal}} = \hat{L}_i^{\text{CORAL}} + b_s \tag{2}$$

The calibration channels were excluded from performance evaluation, leaving 10 test channels from two devices in each target group.

[2] Nowacki B, Li T, Navidi S, Mundiwala M, Hua H, Mai T, Craine D, Miao F, Cooper R and Hu C. ModularML: A backend-agnostic, declarative framework for reproducible and modular machine learning experiments. https://github.com/REIL-UConn/modular-ml.
